# Ending AIDS: trends in the incidence of HIV in eastern and southern Africa

**DOI:** 10.1101/383372

**Authors:** Brian G. Williams, Reuben Granich

## Abstract

While great progress has been made in the control and management of the HIV epidemic there is still much to be done. Using trends in the rate of new HIV infections in eastern and southern Africa we assess the current state of the epidemic and evaluate the future prospects for controlling it. If we let an incidence of 1 per 1,000 people represent a *control threshold* then this has been reached, or will probably be reached by 2020, in eastern Africa and is reachable by 2020 in those southern African countries that do not have particularly strong social and economic ties to South Africa if they continue to scale up their treatment programmes. In South Africa and its immediate neighbours Lesotho, Mozambique and Swaziland, the prospects are less certain. These countries are unlikely to reach the control threshold by 2020 but with sufficient political will and commitment to ‘treatment for all’ could do so by 2030.

There are two important *caveats*. First, reaching the control threshold still leaves 35 thousand new infections a year. As the lessons of polio remind us, finding the last few, hard to reach cases will demand more focussed strategies. Second, ending AIDS will not end HIV and about 35 million people will have to be kept on ART for the next 30 to 40 years unless and until a cure is discovered. Even if we assume a modest cost of, say, US$100 per person per year for ART treatment and support, this corresponds to a continuing financial commitment of US$3.5 Bn per year although this is substantially less than the approximately US$ 40 Bn per year currently committed to HIV and AIDS.

## Introduction

We wish to know whether or not the world is on-track to End AIDS by 2030 or if new approaches need to be considered to reach this target. Here we consider the countries of eastern and southern Africa which account for about two-thirds of all those infected with HIV in the world.

The Spectrum/EPP^1^ and the SACEMA/Williams^2^ models have been fitted to the UNAIDS data^1^ for trends in HIV prevalence and ART coverage in each country. However, there is a critical *caveat*. For some countries the UNAIDS estimates of the trends in the prevalence of HIV have changed substantially between 2015 and 2017 as illustrated with data from Swaziland in Figure 1. It is not clear is why the flattening-off of the prevalence around the year 2000, in the data provided in 2014, was replaced by a significant decline in prevalence around the year 2000 followed by a significant increase in the prevalence around the year 2012, in the data provided in 2017. For this discussion we will use the 2017 estimates from UNAIDS as given on the World Bank web-site.^3^

**Figure 1.**
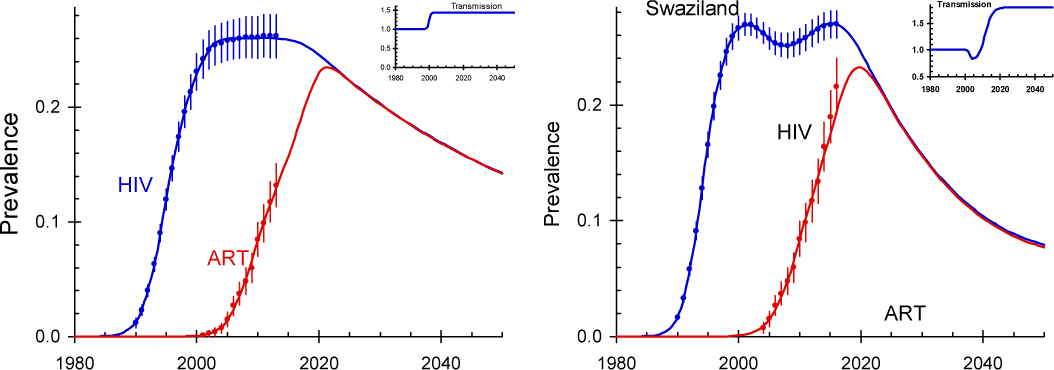
The prevalence of HIV and ART in Swaziland as estimated by UNAIDS in 2014 (left) and in 2017 (right).^1^ The inset labelled ‘transmission’ gives the decrease or increase in the risk of infection arising from changes in behaviour over and above the intrinsic dynamics of the epidemic.^2^

## Methods

With the warning noted above, Figure 2 to Figure 4 give the estimated trends in incidence based on the Spectrum/EPP model (black lines)^4^ and the SACEMA/Williams model (red lines)^2^ fitted to the UNAIDS data. Both models are fitted to the same data for trends in the prevalence of HIV and the coverage of ART. We also show the direct estimates of incidence from the population-based HIV impact assessment (PHIA^5–8^) studies or from other studies using bioassays to estimate incidence, where available^9–13^ (blue dots and lines), and draw lines corresponding to an incidence of 0.1% *p.a.* (green lines) which can be taken as a proxy for epidemic control. The overall trends in incidence from the two models are similar and the PHIA estimates are consistent with the modelled trend data.

**Figure 2.**
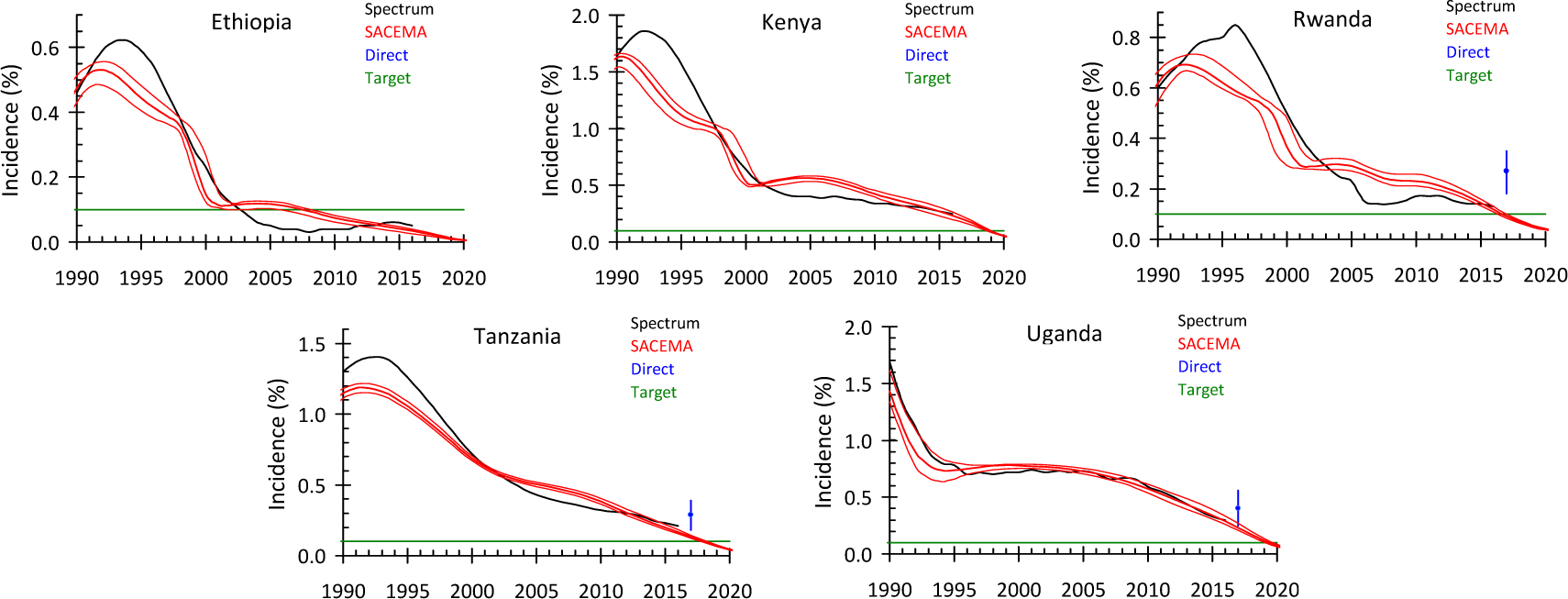
HIV incidence in eastern Africa. *Spectrum* gives the most recent estimates available from the World Bank^3^ which are taken from UNAIDS.^1^ *SACEMA* gives estimates using the SACEMA model^2^ fitted to the most recent time trends in the prevalence and ART coverage made by UNAIDS. *PHIA* are the direct measurements from the PHIA surveys.^6,9,10^ The green lines correspond to an incidence of 1 per 1,000 adults.

**Figure 3.**
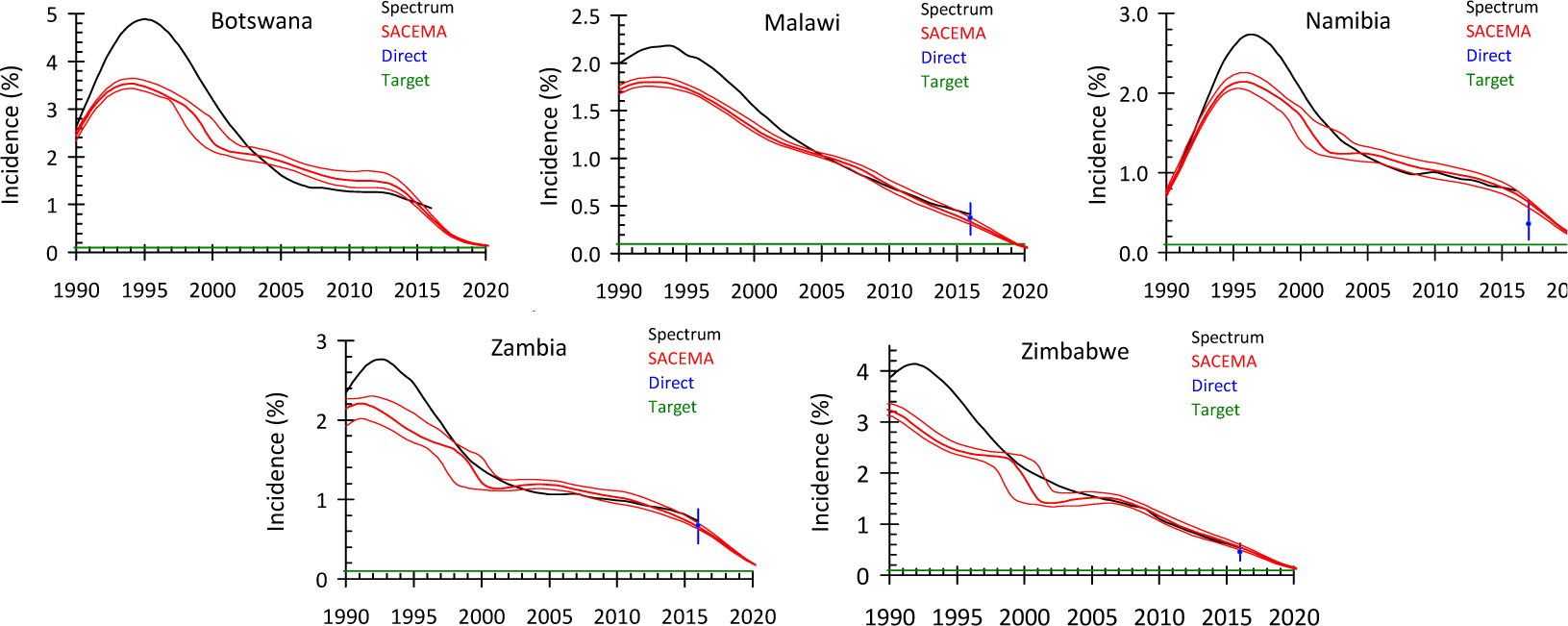
HIV incidence in southern African countries excluding those in Figure 3. *Spectrum* gives the most recent estimates published by the World Bank drawing on the estimates made by UNAIDS.^1^ *SACEMA* gives estimates using the SACEMA model^2^ fitted to the most recent time trends in the prevalence and ART coverage made by UNAIDS. *PHIA* are the direct measurements from PHIA surveys.^7,8,16,17^ The green lines correspond to an incidence of 1 per 1,000 adults.

**Figure 4.**
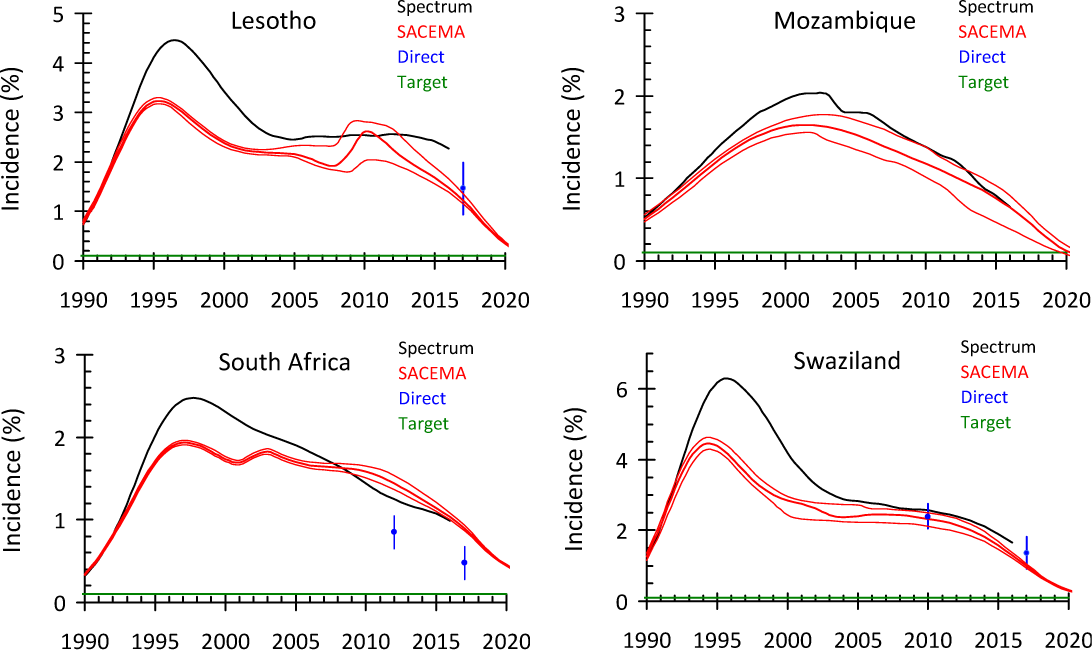
HIV incidence in southern African countries with substantial economic ties to South Africa. *Spectrum* gives the most recent estimates published by the World Bank drawing on the estimates made by UNAIDS.^1^ *SACEMA* gives estimates using the SACEMA model^2^ fitted to the most recent time trends in the prevalence and ART coverage made by UNAIDS. *PHIA* are the direct measurements from the PHIA surveys^18,19^ or from the HSRC surveys in the case of South Africa.^11,13^ The green lines correspond to an incidence of 1 per 1,000 adults.

In the SACEMA/Williams model it is assumed that people at the highest risk are infected before people at lower risk are infected so that transmission parameter, *T*, declines as the prevalence increases as

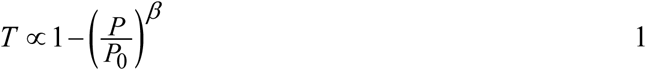

where *P* is the overall prevalence of HIV at any time. 1–*P*_0_ gives the proportion of the population that are not at risk of infection and *β* determines the way in which the transmission parameter converges to the saturation level.^2^

In the Figures below the confidence limits on the SACEMA/Williams model reflect the reported uncertainty in the estimated prevalence of HIV and the coverage of ART while the differences between the two models reflect differences in the model structures.

## Results

In all but two countries the data show a substantial decline in incidence before the year 2000 and therefore before ART became available. In Uganda (Figure 2), the first country in eastern and southern Africa to be affected by HIV, the natural decline in incidence had already occurred by the year 1995 and in South Africa, Swaziland, Lesotho and Mozambique the epidemic started later than in the other countries (Figure 4). This initial decline in incidence is accounted for by the heterogeneity in risk as a function of prevalence. In other words, the initial decline after the early peak is the result of the intrinsic dynamics of the epidemic and changes in the risk behaviour among the populations and does not imply reductions in incidence resulting from treatment of prevention interventions. In the absence of any control interventions one would expect the incidence to fall from the peak to a lower but stable level.

Of greater interest, therefore is what happens after the year 2000 when ART began to be rolled out on a significant scale. By 2005 Ethiopia had reached the control threshold of one new case per thousand adults and Rwanda was not far behind (Figure 2). In Rwanda the direct incidence estimate^10^ is higher than the dynamic models suggest, and this difference needs to be resolved, but with sufficient effort Rwanda could reach the control threshold by 2020.

In Kenya and Uganda there was a period of stasis between 2000 and 2005 but in both countries the incidence declined as ART was rolled out and they are on track to reach the control threshold by 2020 (Figure 2). In Tanzania the incidence of HIV has been in a steady decline since the year 2000 and they are on track to reach the control threshold by 2020.

The epidemic of HIV has been greater in southern Africa than in eastern Africa and all countries in this region are some way off reaching the control threshold. In Botswana (Figure 3), which has a strong HIV control programme, the SACEMA model suggests that they could reach the control threshold by 2020. Malawi (Figure 3) has had a very severe epidemic and is one of the poorest countries in the world but their HIV programme has been a model for other countries in the region^14^ and they are on track to reach the elimination threshold by 2020. In Namibia and Zambia (Figure 3) there has been a slow, steady decline and they will have to scale up treatment more aggressively if they are to reach the control threshold by 2020. Zimbabwe (Figure 3) is an interesting case. The incidence has been falling steadily since the early 1990s and continues to do so.^15^ This is somewhat surprising given the collapse of sensible government over the last twenty years but it is still the case that Zimbabweans are among the best educated in southern Africa and this, combined with a rapid scale up of ART, puts them on track to reach the control threshold by 2020.

The situation in South Africa and the countries most dependent on it is rather worse (Figure 4). Lesotho, which to all intents and purposes is a province of South Africa, had a very severe epidemic, largely as a result of the migrant labour sent to work on the mines in South Africa,^20,21^ and has had a high incidence for the last ten years. The SACEMA model suggests that they are bringing the incidence down and could reach the control threshold by 2025 but if they aggressively scale up treatment. The epidemic of HIV in Mozambique started later and was less severe than in Swaziland (Figure 4) but the incidence has been in decline for the last ten years and they too could reach the control threshold by about 2025 if they aggressively scale up treatment. In South Africa the decline in incidence has been steady but the country is still some way from the control threshold and both models suggest that this will only be reached by about 2030. Swaziland had a very severe epidemic, there was a period of stasis between 2005 and 2012 but the rapid scale up of ART suggests that they could meet the control threshold by about 2025.

It is important to note that the incidence assays are in reasonable agreement with the estimates from the dynamical models. In the countries of eastern Africa, the direct estimates tend to be higher than the modelled estimates (Figure 2) while in southern Africa the direct estimates are in good agreement with the modelled estimates (Figure 3 and Figure 4). An important exception is South Africa where the direct estimates are about two-thirds of the modelled estimates.

## Conclusion

In eastern and southern Africa, the natural dynamics of HIV combined with behaviour change in some countries led to substantial declines in the incidence of HIV before ART became available. After the year 2000 the incidence levelled off in most countries but widespread treatment with ART has brought the incidence down further. Some countries in eastern Africa have reached the control threshold of one case per thousand adults, some in southern Africa are on track to reach the control threshold by 2020 if the roll out of ART continues to expand while South Africa and its immediate neighbours are only likely to reach the control threshold by 2030 and then only if they aggressively expand their treatment programmes. The PHIA data and the HSRC data, using recent infection assays, are generally in good agreement with the model estimates. In Rwanda the direct estimate is higher than the models suggest and in South Africa the direct estimate is lower than the models suggest.

Finding and treating the last 0.1% of the population that is infected each year and making sure that others are not infected will remain a challenge and will demand more focussed and imaginative approaches to case finding, treatment and prevention for those that remain at risk. While ending AIDS by 2030 can be achieved by 2030, ending HIV will depend on maintaining the roughly 35 million people currently infected with HIV on treatment for the next 30 to 40 years, when they will die of natural causes not related to HIV, unless and until a cure can be developed.

